# Automated Viability Estimation from Digital Holographic Microscopy: Validation on Heterogeneous Industrial Bioproduction Cultures

**DOI:** 10.64898/2026.03.10.710837

**Authors:** Guillaume Godefroy, Anais Berger, Eric Calvosa, Tigrane Cantat-Moltrecht, Gaétan Girard, Emmanuel Guedon, Lionel Hervé, Angéla La, Thomas Saillard, Stanislas Lhomme

## Abstract

Cell viability is a critical parameter in bioproduction, yet most facilities still rely on manual, offline assays. This work introduces a new label-free Digital Holographic Microscopy (DHM)-based viability prediction pipeline using a simple optical design compatible with both on-line and in-line probe implementation. Unlike previous approaches validated under controled laboratory conditions, the proposed pipeline was designed to operate across diverse CHO bioprocess conditions without calibration or parameter tuning. It was validated on a large, heterogeneous dataset comprising 40 cell cultures collected from industrial and academic sites, spanning multiple cell lines, culture media, process modes and cell densities up to 100 million cells/mL. Beyond viability estimation, exploratory analyses suggest that DHM-based monitoring can provide additional process-relevant insights, including early detection of viability decline and correlation with recombinant protein titer. Together, these results indicate that DHM has the potential to enable a new generation of non-invasive, multiparametric monitoring tools for advanced bioproduction control.

## Introduction

Biomedicines are increasingly present in the therapeutic arsenal available to healthcare professionals. Among these products, monoclonal antibodies and recombinant proteins produced in mammalian cells such as Chinese hamster ovary (CHO) cells are among the most prominent^1^. Producing these molecules remains costly due to high material expenses, labor-intensive operations, and the need for strict quality assurance^2–4^. One promising path to lowering these costs is to improve the quality control chain to boost culture productivity by increasing automation, enhancing process monitoring, and reducing manual interventions^5^.

One of the most critical parameters to monitor in any bioproduction process is cell viability. Drops in viability can indicate early signs of contamination, nutrient depletion, or shear stress, which, if undetected, can lead to reduced yield or batch failure^6^. It serves as a criterion for determining the feeding rate, the time of induction or transfection, and the harvest point^7^. It therefore directly impacts culture productivity and product quality. Yet, to monitor viability, most facilities still rely on manual, offline assays, such as trypan blue exclusion^8^ or fluorescent dyes combined with flow cytometry^9^. These methods require sampling, dilution, and staining with toxic dyes; steps that introduce contamination risks^10^, operator-dependent variability^11^, and incur material and labor costs, as well as sample loss. These constraints limit the temporal resolution of the measurements, which are usually spaced by 12 to 24 hours, allowing critical events such as apoptosis, nutrient depletion, or contamination to go unnoticed for hours. In response, industry is moving toward automatic approaches compatible with in-line and on-line monitoring. In-line refers to probes placed directly inside the bioreactor, while on-line refers to probes connected to the bioreactor via external loop.

Significant advances have been made in developing alternative automatic label-free approaches compatible with in-line and on-line implementation. For example, culture optical density, which correlates with viable cell concentration, can be measured using back-scattered light turbidity sensors^12^, while biocapacitance probes exploit the dielectric properties of viable cells with intact membranes^13^. Spectroscopic approaches, including UV/Vis, IR, Raman, and even NMR, have also been explored for bioprocess monitoring^14^. However, for all these techniques, viability is derived indirectly and remains sensitive to changes in cell morphology, accumulation of non-viable or lysed cells, debris, and environmental fluctuations, requiring complex modeling and extensive calibration to ensure accuracy^6,15,16^.

Imaging-based methods have therefore emerged as attractive alternatives, as they enable single-cell–level viability assessment. Two-photon fluorescence lifetime imaging microscopy (2P-FLIM) has recently been proposed, inferring viability from the autofluorescence of intracellular metabolites^17^, but the complexity of its optical setup significantly limits its potential for in-line deployment. In contrast, bright-field imaging relies on a simple optical configuration and has already been implemented for direct monitoring inside bioreactors^18,19^. Although morphological features observable in bright-field images have been shown to correlate with cell viability^20^, the method’s reliance on intensity images alone can be limiting, given the high transparency of cells. Moving to Digital Holographic Microscopy (DHM) can offer substantial advantages. While preserving the possibility of using a simple optical setup, DHM reconstructs not only intensity, but also the optical phase shift induced by cells. This phase information correlates with cell dry mass, thickness, and refractive index, which are fundamental biophysical parameters for cell characterization^21,22^. DHM therefore shows strong potential to extend bioprocess monitoring beyond cell viability measurement. Accordingly, prior studies have demonstrated that DHM can discriminate apoptosis from necrosis^23–25^, can identify infected versus uninfected cells^26^ and can even predict adeno-associated virus productivity in Sf9 cultures^27^. These capabilities position DHM as a promising tool for next-generation bioprocess monitoring.

To date, viability estimation approaches based on DHM^23,24,28,29^, as well as bright-field imaging, have been validated under controlled and homogeneous conditions. None of these studies aimed to establish a general algorithm applicable across variable culture conditions and process configurations. In contrast, industrial bioproduction environments exhibit substantial variability, encompassing diverse CHO cell lines, media formulations, and operation strategies, and the field continues to evolve rapidly^30,31^. Such variability can significantly affect cell morphology, image contrast, and viability dynamics throughout the culture. Moreover, most existing in-line compatible viability methods have been developed and validated only at low to moderate cell-density conditions, whereas high-density cultures are gaining increasing interest in the field as a strategy to improve productivity. These dense environments, exceeding 100 million cells/mL^32,33^, introduce challenges for all techniques: at the time of writing, no method has reliably demonstrated viability monitoring at such high cell densities. Optical density and capacitance sensors suffer from signal saturation and poor sensitivity to viability changes, while bright-field and DHM imaging encounter overlapping cells and reduced contrast^34^.

This work addresses several of the challenges described above.

Its principal contribution lies in tackling the development of a generalizable pipeline for cell viability measurement across a wide range of culture environments, without requiring additional calibration or parameter tuning. This is achieved through a DHM-based imaging approach combined with a novel viability estimation algorithm, validated on a uniquely large and heterogeneous dataset collected from three industrial production sites and one academic laboratory. The dataset encompasses multiple CHO cell lines expressing monoclonal antibodies, diverse media formulations, and varied production processes. The study also addresses the challenge of operating under very high cell-density conditions, reaching approximately 100 million cells/mL, by optimizing both the image acquisition strategy and the reconstruction process. in addition, the work explores the potential of DHM imaging beyond viability estimation through exploratory results on early anticipation of viability declines and on the prediction of recombinant protein titer.

This study was conducted using an offline optical bench to optimize data collection. However, with the ultimate objective of enabling on-line or in-line deployment, DHM was implemented using a defocused transmission microscope configuration in which the beam interferes with itself. Compared to conventional DHM systems relying on interferometric setups with a dedicated reference arm^35^, this configuration substantially simplifies the optical design and reduces cost.

The paper first describes the image acquisition, reconstruction, and segmentation procedures. An analysis of the dataset is then presented to highlight its heterogeneity and, consequently, the challenge of developing a generalizable viability measurement algorithm. The viability measurement algorithm is subsequently detailed, followed by results addressing the three objectives outlined above.

### Acquisition, reconstruction and image segmentation

#### Image acquisition

The image acquisition and pre-processing pipeline is described in Figure 1. The process starts with image acquisition, followed by phase and absorption image reconstruction, image segmentation, and single-cell metrics extraction. DHM is a label-free imaging technique that records the interference pattern generated by the sample under study^36^. Quantitative phase and absorption images are then computationally reconstructed from this pattern. The phase component represents the optical phase delay introduced by the sample, which reflects variations in optical path length caused by differences in refractive index and thickness. For clarity, this component is expressed as the optical path difference (OPD, in *µ*m) and referred to as the “phase image” throughout the manuscript. The amplitude component is dimensionless and proportional to the local light attenuation, which includes contributions from both absorption and scattering. In this study, this component is referred to as the “absorption image” throughout the manuscript. The absorption images are normalized to the background and expressed as percent transmission (%T).

**Figure 1.**
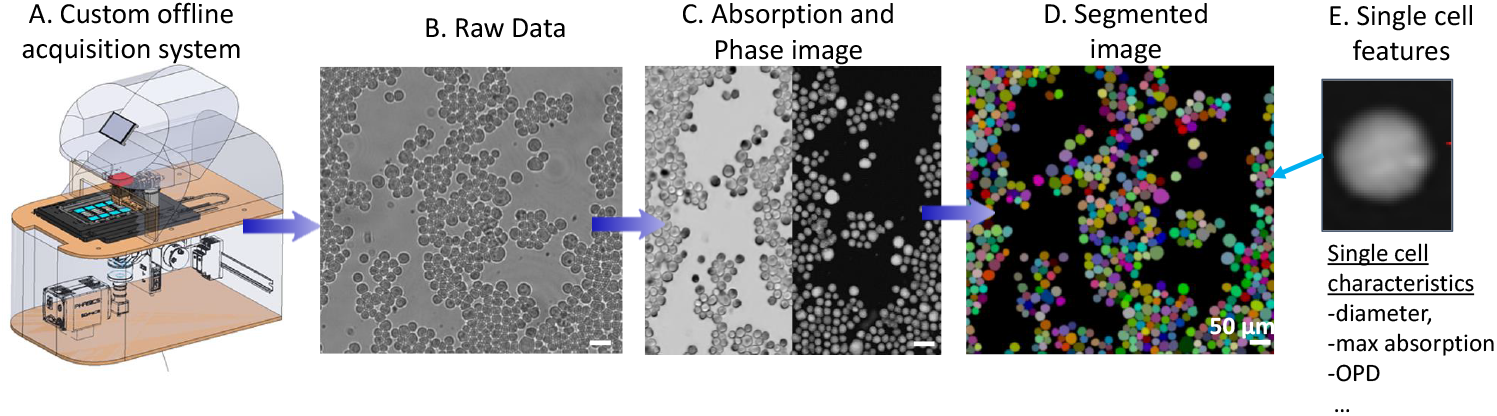
Image acquisition and pre-processing pipeline. Holograms are acquired using the DHM microscope, after which absorption and phase images are reconstructed. A segmentation algorithm is then applied to identify individual cells and single-cell features are extracted.

DHM was implemented through a defocused transmission microscope configuration, in which the optical beam interferes with itself. To balance spatial resolution, robustness, and field of view, a low-numerical-aperture (0.25 NA) air objective was employed. Unlike some other DHM systems that employ higher NA objectives to enhance morphological detail, our configuration provides a sufficiently large field of view to image a representative number of cells per acquisition while maintaining a compact and easily adaptable setup. For the purpose of this study, to enable automated acquisition of a large dataset, the DHM was integrated into an in-house-designed optical bench equipped with a motorized scanning stage, allowing for rapid sequential image acquisition (Figure 1A). Additional details are provided in the Materials and Methods (Section A).

#### Image reconstruction and segmentation

Phase and absorption images were reconstructed using an updated version of the algorithm described in^37^. The reconstruction process involved alternating between two approaches: inverse-problem optimization and artificial neural network inference. The optimization started with several iterations of a conventional gradient descent optimization algorithm, resulting in a preliminary estimate of the sample. This image was then provided to a convolutional neural network, referred to as the reconstruction CNN throughout the manuscript. This neural network aids in reducing phase wrapping errors and improves reconstruction resolution, particularly in cases of high cell density. The predicted image served as the initialization of a second and final reconstruction step, which corrects to a certain extent the neural network prediction errors and re-anchors the reconstructed images to the measurement data. As described in Materials and Methods (Section B), significant improvements were introduced. These include the use of two different neural network models for phase and absorption image reconstruction, the refinement of the simulation dataset for phase reconstruction, and the use of an experimental dataset for the absorption reconstruction.

In the reconstructed images, individual cells were identified and segmented using the Cellpose framework^38^. Cellpose is a generalist deep-learning–based segmentation algorithm that generates a unique mask per cell. The model was fine-tuned to match our image conditions.

### Assessing cell viability across culture conditions

#### Culture variability

To address the challenge of developing a generalizable pipeline, images of cells cultured under a wide range of conditions were collected through experiments conducted in both research and industrial environments. Experiments involved monitoring a CHO cell culture over one to two weeks, covering an initial growth phase with high viability followed by a decline phase, encompassing a broad spectrum of physiological states and temporal dynamics of growth, production, and cell death.

The dataset encompassed six CHO cell lines (described in Materials and Methods, Section i), which exhibited distinct morphological behaviors. For example, one cell line showed a marked increase in cell size over the course of the culture, whereas another maintained a relatively constant size. Five different culture media were used, introducing variability in refractive index contrast and consequently in the quantitative phase signal, since this signal depends on the refractive index difference between the cell and its surrounding medium. Cultures were conducted under three process modes (batch, fed-batch, and perfusion), which impacted the cells’ morphological and phase features. For instance, cells cultivated under fed-batch conditions displayed a consistent and pronounced increase in optical path difference (OPD) over time, in contrast with those grown in batch mode. In a few cases, apoptosis was triggered by nutrient exhaustion (feeding stop) resulting sudden metabolic stress. Experiments were performed across several culture vessels, including bioreactors of various capacities (up to 50 L), Ambr bioreactor systems, shake flasks and spin tubes, with additional variations in feed composition, pH adjustment strategies, and the use of antifoam agents. A table summarizing all encountered conditions and describing cell lines is provided in the Supplementary Materials. Cell line names and media compositions have been anonymized for confidentiality reasons.

All these conditions resulted in a wide variety of viability dynamics, as shown in Figure 2. In some cases, viability dropped abruptly, while in others it declines gradually. Cultures lasted only a few days or extend over two weeks.

**Figure 2.**
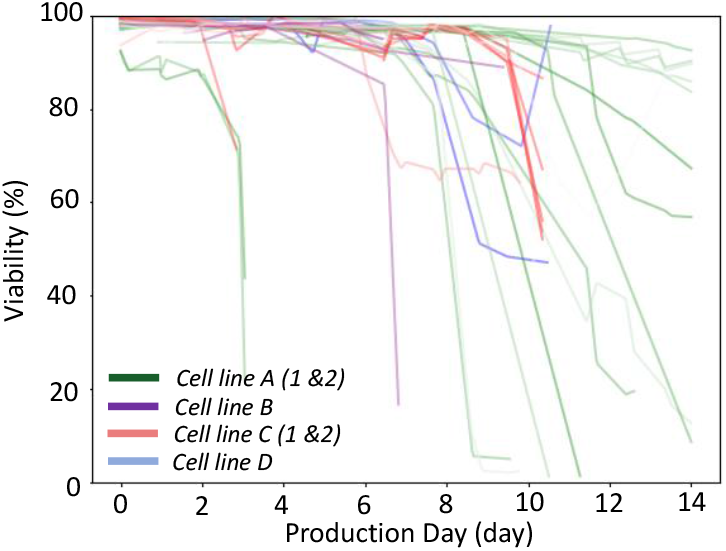
Viability evolution of the 40 cultures analyzed in the study, colored by cell line.

#### Individual cell features are insufficient to measure viability

Figure 3 illustrates the variability observed both between cultures and within individual cultures. Part A shows images of representative viable and non-viable CHO cells from three different cell lines, acquired with our microscope in both the DHM modality (reconstructed as a phase image) and a more conventional bright-field modality (simple transmission microscopy at focus). Each cell is imaged using both modalities quasi-simultaneously (within an interval of < 1 s). Viable cells from cell lines A.1 and D display higher granularity in the absorption image compared to cells from cell line B. Phase values also vary substantially across cell lines. In comparison, non-viable cells exhibit lower OPD and absorption values, and increased granularity. However, clear thresholds distinguishing viable from non-viable cells seem difficult to define based solely on these images.

**Figure 3.**
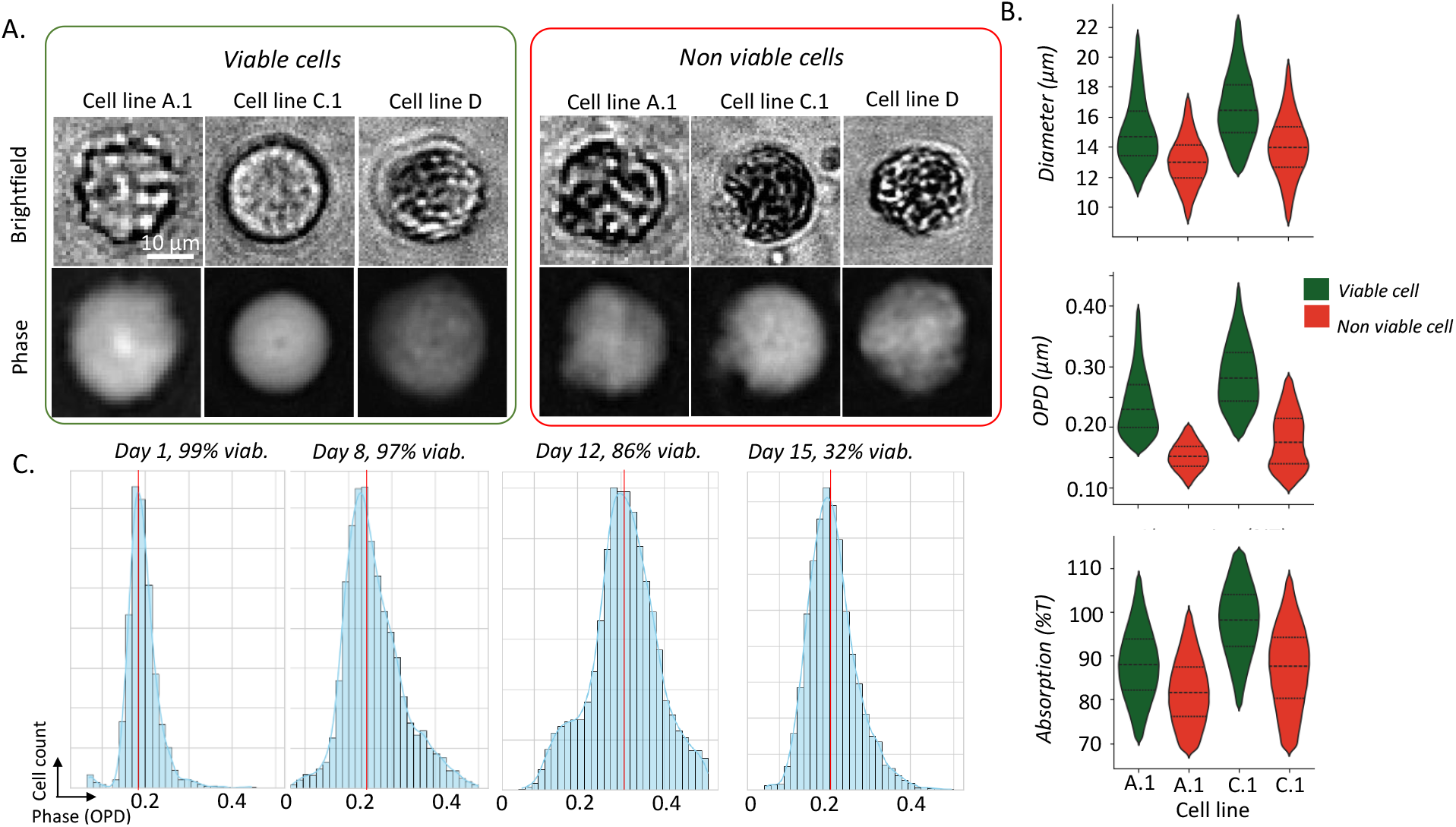
Illustration of the heterogeneity across CHO cell lines. (A) Representative phase and bright-field images of three CHO cell lines. (B) Violin plots showing intra- and inter-cell line variability of quantitative and morphological features. (C) Visualization of intra-culture variability in the phase signal, illustrated by the distribution of the phase component at four time points during the culture.

Part B highlights the strong heterogeneity across and within cell line populations, focusing on three cell features: cell diameter (*µ*m), mean OPD (*µ*m) over the cell area (*µ*m^2^) and mean absorption around the cell’s center of mass (%T). Among a single cell line, the overlap between viable and non-viable cells is already significant (Figure 3B, first graph). The complexity increases when considering multiple cell lines. For instance, the absorption distributions of viable cells from line A.1 and non-viable cells from line B almost completely overlap (Figure 3B, third graph). Such heterogeneity can be partially explained by differences in culture conditions represented in these distributions, but intrinsic variability between cell lines remains an important source of variation.

In panel C, intra-experimental variability is examined by analyzing the temporal evolution of OPD values in an experiment conducted with cells from cell line A.1. Four OPD histograms computed at different time points of the culture are presented. For day 1 and day 8, although viability remains similar, an increase in OPD is observed. At day 15, when most cells are dead and viability drops to 32%, the mean OPD value is close to that of day 1, which corresponds to a population of mostly viable cells.These observations indicate that simple thresholding of feature values is insufficient to accurately determine cell viability.

#### The population analysis algorithm

These observations motivated us to design our algorithm around a population-based multiparametric analysis rather than relying solely on individual cell features. The algorithm was explicitly designed to operate independently at each time point, without requiring information from earlier or later culture states. This choice ensures that the proposed approach, and any future device implementing it, can function in a fully stand-alone manner.

The principle of the analysis at a given time point is illustrated in Figure 4. Two single-cell features are first considered: the mean OPD over the cell area and the mean absorption around the cell’s center of mass. A 2D histogram of these two features is constructed, representing the distribution of cells at this time pointin the OPD–absorption space. The number of distinct cell populations present in the culture in then determined through a local maxima detection algorithm^39^ applied to the histogram, treated as an image, yielding either one or two populations.

**Figure 4.**
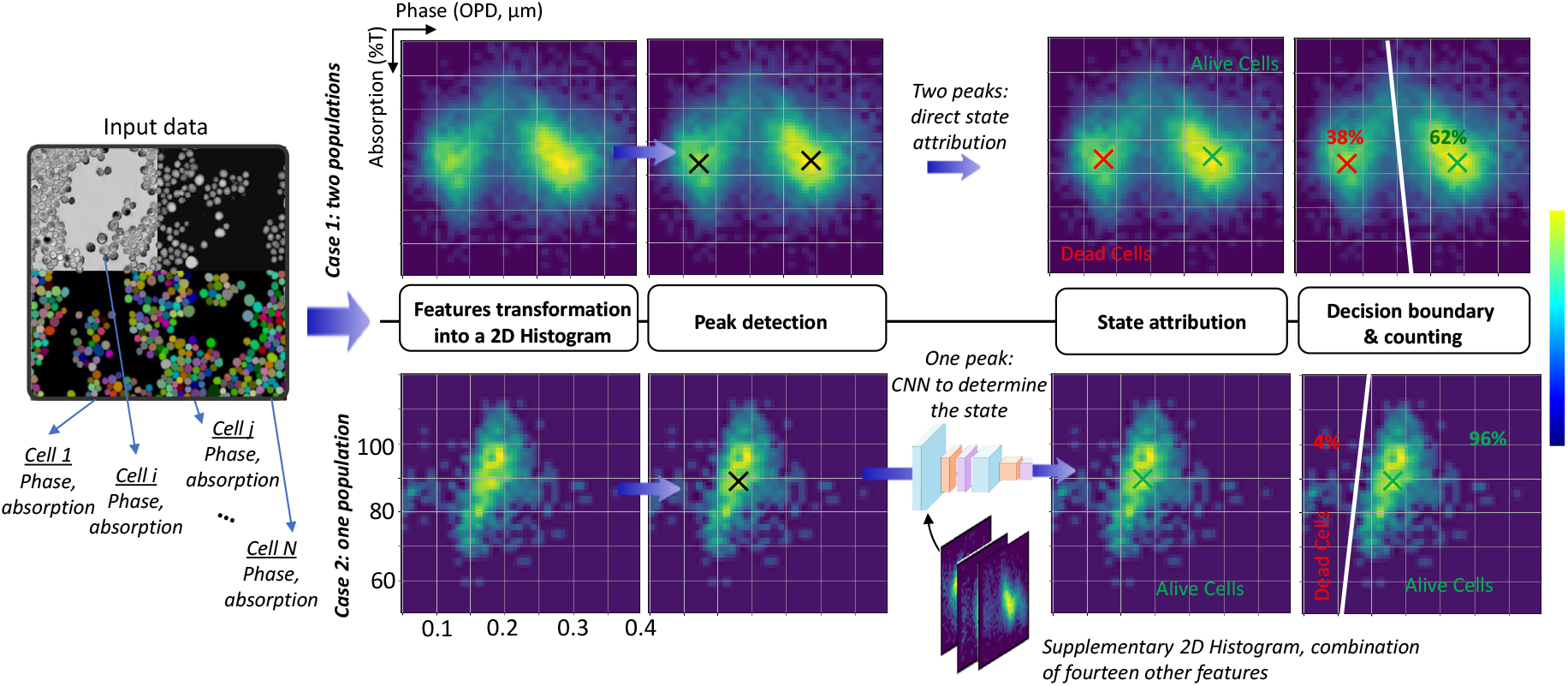
The viability measurement algorithm. Single-cell features are extracted and transformed into a 2D histogram. Peak analysis is then performed: if a single peak is detected, a trained CNN classifies the population state; if multiple peaks are present, cells with lower phase values are classified as non-viable and those with higher values as viable. Predefined rules then define the decision boundary to estimate overall culture viability.

At a given time point, non-viable cells consistently exhibit lower OPD values than viable cells. Therefore, when two populations are detected (case 1 in Figure 4), a decision boundary is drawn with a fixed slope and positioned relative to the centers of the two populations. Cells are then counted on each side of this boundary to compute viability. When only one population is detected (case 2 in Figure 4), an additional classification step is performed to determine whether the culture predominantly contains viable or non-viable cells. As shown previously, it is not straightforward to define a simple rule based on distribution moments, since the features of viable and non-viable cell populations often overlap, even within the same culture between different time points. To overcome this limitation, a convolutional neural network, referred to as the prediction CNN throughout the manuscript, was trained to predict the dominant viability state of the population.

The network takes as input not only the OPD/absorption histogram but also nine additional 2D histograms constructed from combinations of ten other single-cell features. Indeed, phase and absorption alone are not always sufficient to distinguish between the two states, as illustrated in Figure 3C. Once the dominant population has been identified, a parameterized decision boundary is applied to separate the space, selected according to the CNN prediction. Details about the features selection and the training procedure are available in Materials and Methods (Section E).

For validation and subsequent analyses, the prediction CNN was trained using a one-vs-all approach in which one culture was reserved as the test set, and all others were used for training and validation.

#### Results on the overall dataset

The prediction CNN consistently identified the correct dominant population state based on the histograms, which was a prerequisite for applying the viability prediction pipeline. Indeed, any error made by the prediction CNN would lead to a complete misanalysis of the situation, since it determines on which side of the population the decision boundary is placed. The other parameters of this linear boundary, its slope and precise position relative to the population(s), were carefully optimized to maximize viability prediction accuracy across the 40 cultures.

As a ground truth viability estimation, a Beckman Coulter Vi-CELL (Beckman Coulter, Roissy, France) was used. Table 1 summarizes the results computed over 550 time-point measurements, showing a high level of agreement at elevated viability levels, with a deviation of 1.8% ± 1.5% for viability values between 90% and 100%. The spread increased at lower viabilities, but remain limited to 3.7% ± 5.7% over the entire viability range.

**Table 1.** Comparison of viability estimates with Vi-CELL measurements.

| Viability range | Error mean | Error std | Error max |
| --- | --- | --- | --- |
| 95 – 100% | 1.6% | $\pm 1.4\%$ | 8.5% |
| 90 – 100% | 1.8% | $\pm 1.5\%$ | 8.5% |
| 70 – 100% | 2.4% | $\pm 3.4\%$ | 13.7% |
| 0 – 100% | 3.7% | $\pm 5.7\%$ | 49.5% |

These results are highly encouraging. For industrial applications, the primary objectives are to maintain a high level of viability and to detect the starting point of culture decline, in order to determine when to trigger harvest, tipically above 70%^5–7^. Our method shows good accuracy within this critical viability range. As illustrated in Figure 5 (right column), the predicted viability closely follows the Vi-CELL measurements, particularly above 90%, and the initial drops in viability were detected simultaneously by both methods in those three representative cultures. The culture shown in Figure 5 (cell line D) was deliberately selected because these criteria are met even when substantial deviations from Vi-CELL measurements occur at low viability. It is important to note that deviations are expected between the Vi-CELL measurements and our viability estimates, as the two systems rely on different criteria to assess viability. Consequently, the definitions of “viable” and “non-viable” cells differ between the two methods. The Vi-CELL, which relies on trypan blue staining, classifies cells according to membrane permeability, thereby reporting loss of membrane integrity. In contrast, DHM captures morphological and quantitative biophysical changes. As a result, discrepancies between the two measurements are particularly expected during the transition from viable to non-viable states. It should also be noted that four different Vi-CELL instruments were used, each with its own custom setup. Interestingly, flow cytometer analysis for Experiment B (Figure 5 cell line A.1, yellow curve), reveals discrepancies with the Vi-CELL measurements (approximately 3% on average for this experiment), even though both methods are considered gold standards for assessing viability. This further underscores that the estimated viability can vary slightly, depending on the criteria chosen to distinguish viable from non-viable cells.

**Figure 5.**
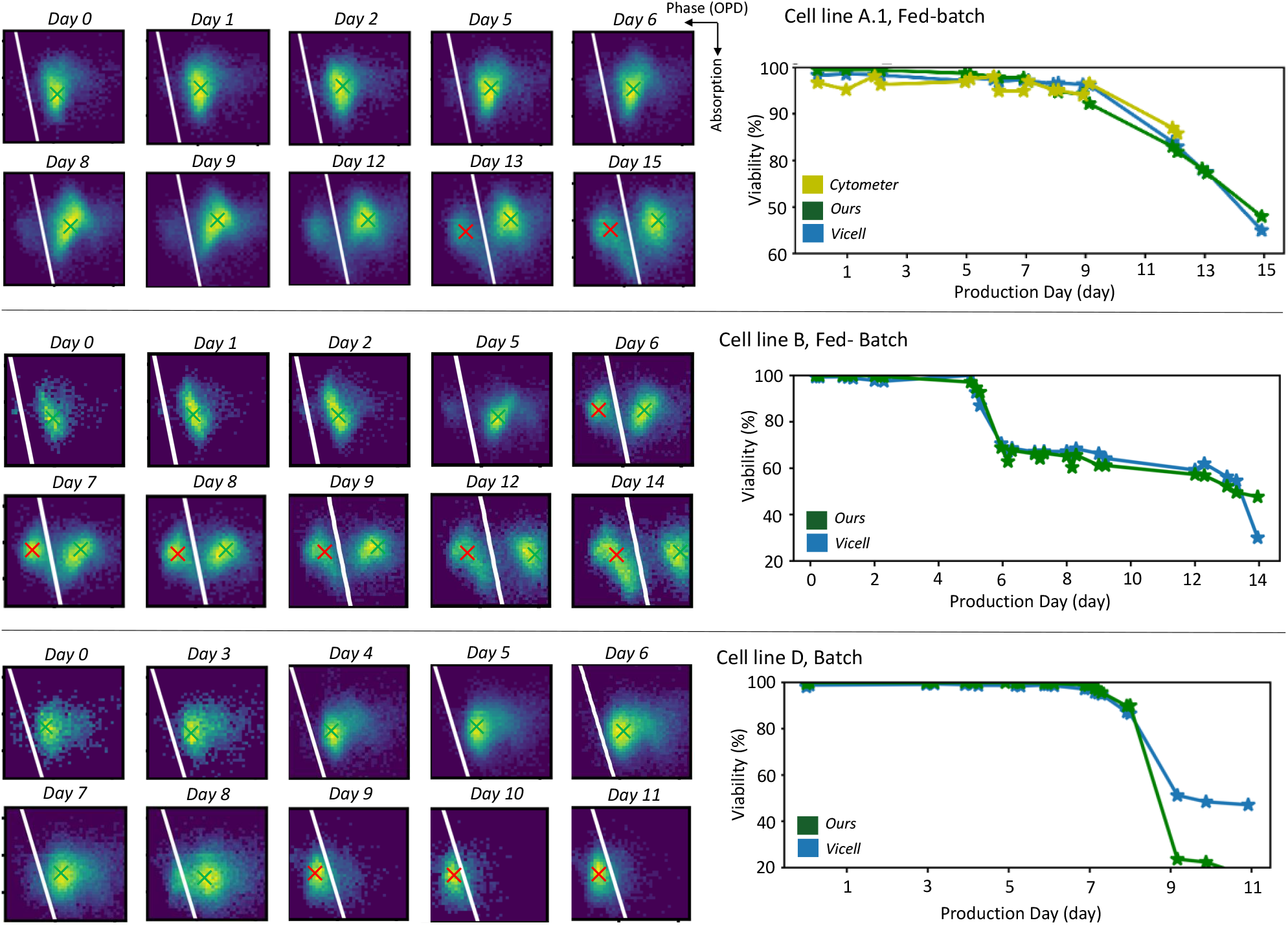
Application of the DHM-based viability prediction pipeline along three culture processes: evolution of 2D histograms (a) and predicted viability compared with Vi-CELL measurement (b).

#### Focus on high densities

Achieving high-density cultures, reaching up to 100 × 10^6^ cells/mL, represents a major milestone for the biopharmaceutical industry. At such densities, cells become highly confined, and diffraction patterns rapidly overlap, making phase and absorption image reconstruction particularly challenging (Figure 6A). The entire pipeline was designed from the outset to address these challenges to accurately handle images acquired under high-density conditions. The reconstruction CNN used in our previous work^37^ was adapted, and the defocus distance (the offset between the objective focal plane and the sample plane) was carefully optimized and deliberately kept small (50 *µ*m) to minimize fringe overlap between neighboring cells (details in Materials and Methods, Sections A and B).

**Figure 6.**
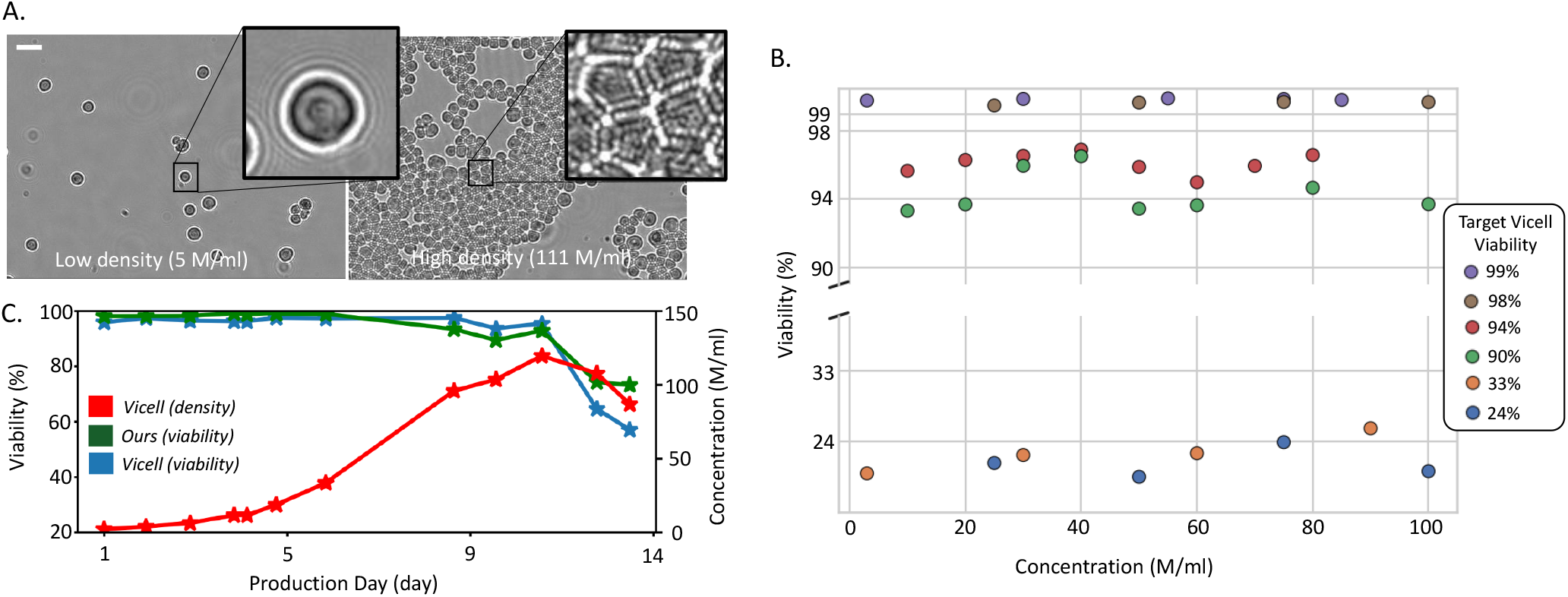
High-density results. (A) Illustration of diffraction-pattern overlap observed at high cell density. (B) Viability estimation on manually mixed populations. (C) Monitoring of a perfusion bioreactor culture.

The pipeline was first applied to manually mixed populations of viable and non-viable cells, i.e., artificial samples generated with controlled viability and density. The measured viabilities, corresponding to reference values of 99%, 98%, 94%, 90%, 33%, and 24%, are shown in Figure 6B for different cell densities. The deviation between our method and the reference showed no significant difference between low- and high-density samples, ranging from from 0-6% at high viability and 2-12% at low viability. The average error was 5% for concentrations below 50 × 10^6^ cells/mL and 5.5% above this threshold. To further evaluate performance under real high-density conditions, the pipeline was applied to a perfusion culture reaching 110 × 10^6^ cells/mL (Figure 6C). The viability measurements closely matched those obtained with a Vi-CELL, both in absolute values and in identifying the onset of viability decline. While these results are encouraging, additional experiments under varied conditions are required to fully validate performance at such very high cell densities.

### Exploratory results: going beyond viability measurement

#### Estimating recombinant protein titer during culture

As an imaging technique, DHM offers the key advantage of directly observing cells, unlike indirect methods such as biocapacitance or turbidity sensors. This capability opens the door to monitoring additional parameters, as demonstrated in the study by Pais et al.^27^, in which the authors successfully inferred AAV titers from DHM images of insect cell cultures.

Inspired by this study, we investigated whether our extracted features could be correlated with bioreactor production output, represented here by the titer of a recombinant protein, the immunoglobulin G (igG). The ten single-cell features were extracted, and their distributions were evaluated for their ability to model IgG levels. Histograms were computed for each feature and used as inputs to a gradient boosting regression model trained to predict the measured IgG titer. A one-versus-all training/testing strategy was applied to ensure sufficient variability in the training set and independent evaluation on multiple cultures (details in Materials and Methods, Section H). Test-set predictions are shown in Figure 7A. In all five illustrated cases, the predicted trajectories successfully captured the experimental trends: when IgG levels increased, the model predictions increased simultaneously, and when no increase occurred, the predicted values remained low. Some discrepancies in absolute values were observed, particularly for culture (i), but the overall dynamics were well reproduced. Mean errors for different IgG concentration ranges are summarized in Figure 7A (right panel). These results are encouraging, as they suggest that holographic features contain exploitable information related to IgG titer. Further work is required to determine whether these features can be used for quantitative prediction across a broader range of conditions, or whether the model’s applicability is restricted to experiments closely matching those used for training.

**Figure 7.**
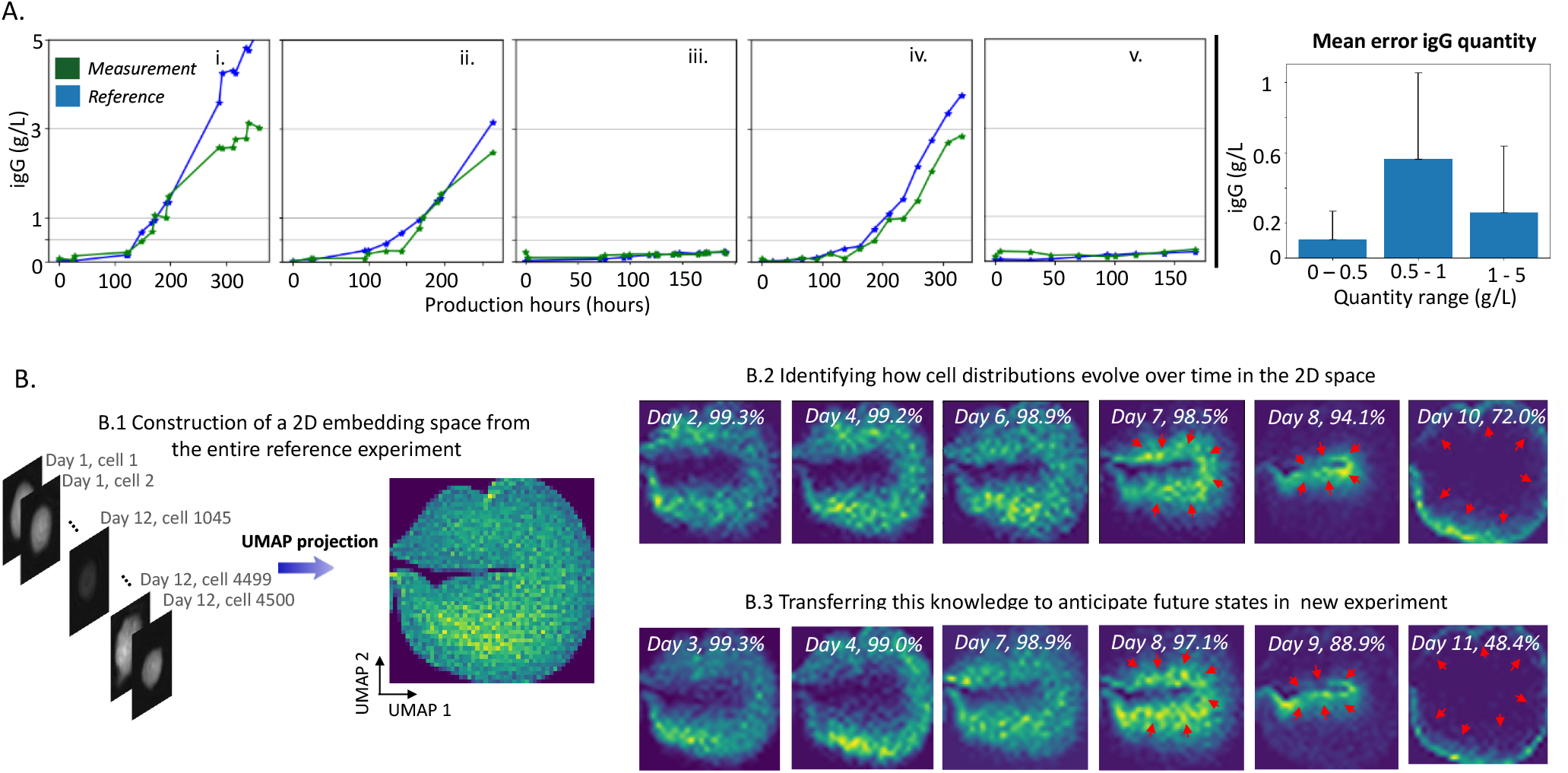
(A) Estimation of IgG production during culture. Five IgG production profiles are shown, along with the average prediction error across different IgG concentration ranges. (B) Anticipation of culture-state transitions using a UMAP projection. The UMAP space is constructed using all data from a training experiment (B.1). The temporal evolution of cell distributions within this space is shown for the training experiment (B.2) and subsequently for an independent test experiment (B.3)

#### Anticipating culture-state transitions, such as the onset of viability decline

When examining the temporal evolution of histograms and other extracted metrics, we observed that several features begin to change before the estimated viability decreases. For example, consistent with a previous report^24^, some experiments exhibited an increase in refractive index prior to the onset of viability decline, potentially reflecting early apoptotic events. This motivated us to investigate whether the temporal evolution of cell features could be modeled and used to detect early changes. We focused on two comparable experiments (same cell line and same culture process), one serving as the reference to define the model, and the other as the test case. Since individual metrics alone were insufficient to capture such subtle behaviors (for example, the refractive index increase was not consistently observed), a multivariate approach was employed.

The method and results are illustrated in Figure 7B and detailed in Materials and Methods (Section G). Briefly, the ten single-cell features from all time points of the reference culture were embedded using UMAP^40^, which projects high-dimensional data into a 2D space (7B.1). This embedding provides a representation that captures the entire range of cellular states encountered throughout the culture. We then examined how the cell population moved through this space over time. Up to day 7, the population distribution did not exhibit substantial changes and approximately uniformly covered the entire structure of the projection. From day 7 to day 8, however, the cells progressively shifted toward the center of the space, coinciding with the onset of viability decline (Figure 7B.2). By day 10, when viability had dropped significantly, the population had migrated toward the outer rim of the embedding. Detecting this initial inward movement could therefore provide an early warning signal that the culture is approaching viability decline. Next, cells from the second, unseen culture (test experiment) were projected into the same UMAP space, time point by time point, as would be done in continuous monitoring (Figure 7B.3). The temporal trajectory of the cell population was highly similar. This suggests that, for experiments performed under comparable conditions, population trajectories in UMAP space are reproducible and could potentially be monitored on-the-fly to trigger alerts when the culture enters a predefined transition zone.

These results remain exploratory and are currently restricted to cultures with similar parameters. Additional work is required to assess how well the reference model generalizes to experiments conducted under different conditions (e.g., different media, feeding strategies, or cell lines). Moreover, the design of an alert-generation algorithm based on these trajectories must be formalized and made easily adaptable when changing the reference culture.

## Discussion

Overall, the results indicate that our DHM-based viability measurement pipeline has strong potential for broad applicability. The diversity of the dataset suggests that the algorithm generalizes across a wide range of industrial CHO cultures without requiring modification. Because some decision parameters (e.g., the position of the decision boundary) are engineered, the method also remains adaptable: if necessary, these parameters can be fine-tuned to further improve alignment with Vi-CELL measurements in specific settings.

The use of the prediction CNN represents a double-edged sword. On the one hand, it enables the application of the same algorithm uniformly across all experiments. On the other hand, it introduces additional complexity, and if the CNN fails on a new type of culture, retraining would be required, a process that is both costly and time-consuming. This is why we also developed a fully handcrafted version of the algorithm, relying solely on rule-based histogram classification to determine whether the population is composed mainly of viable or non-viable cells, which could therefore replace the prediction CNN. Comparable results were obtained, but at the cost of cell-line specific adjustments.

While this study focused on CHO cells cultivated in suspension, we believe that the algorithm and its underlying principles could also be applied to other cell types, such as HEK or insect cells, and potentially even lymphocytes, provided that appropriate optical adaptations are made to accommodate their smaller size. Adherent cell cultures could also be considered, as they exhibit pronounced morphological changes during cell death that have been observed with DHM^41^.

We did not discuss concentration measurements in detail. Our segmentation accuracy is consistently around 98% compared with manual annotation (details in Materials and Methods Section C), indicating that cells can be reliably counted in each acquired image. However, the representativeness of this count with respect to the true cell density in the bioreactor lies beyond the scope of this study. In practice, challenges associated with on-line or in-line implementation, such as sensor saturation, non-linear sampling of the cell density within the imaging chamber, or flow-dependent biases^42^, are likely to become the dominant sources of error rather than the segmentation itself. It is worth noting that one advantage of our technique is its speed: acquiring a single image takes less than one second, and analysis typically requires around 30 seconds, depending on the computing hardware and the number of cells per image. For example, in a culture with a cell density of 10^6^ cells/mL, roughly 500 cells can be analyzed per minute. Accumulating images over a few minutes further improves statistical robustness and reduces sampling noise.

Our results on early viability decline detection and titer prediction highlight the potential of DHM to support bioreactor monitoring beyond viability assessment, in contexts where competitive viability-measurement technologies typically cannot operate. These results remain exploratory and require substantial development and validation to determine their practical scope of application. In particular, we do not expect to obtain a single, universal algorithm that generalizes automatically across all industrial processes, as is the case for the viability estimation pipeline. Nevertheless, these advanced features have the potential to provide significant value to industrial users who seek close monitoring of their cultures and are willing to undergo an algorithm-specific optimization phase.

The next step toward industrial application will be the design of an in-line or on-line probe, which presents specific technical challenges. As mentioned earlier, the optical design of our setup is relatively simple, and its miniaturization therefore appears feasible. The primary difficulty in implementing holography within an in-line probe lies in the requirement for a very thin imaging chamber, typically between 20 and 100 *µ*m in depth depending on the maximum cell concentration. If too many cells are present between the light source and the sensor, a usable hologram cannot be formed: diffraction fringes from different cells overlap, and multiple scattering events degrade the holographic signal and even the coherence of the detected light. Dedicated designs will therefore be required to control the imaging thickness while still allowing a sufficient number of cells to enter the chamber. Promising approaches have already been proposed, such as “gated” mechanisms that allow cells to flow into the chamber and then close to maintain a controlled imaging thickness during acquisition^19,43^.

In summary, this work introduces a new label-free viability measurement pipeline based on DHM imaging using a simple and compact optical configuration. The pipeline was validated on a large and diverse dataset including cells from both industrial and academic sites, cultivated under different processes and in various culture vessels. It was successfully tested on high-density cultures, reaching concentrations of up to 100 × 10^6^ cells/mL. Beyond viability assessment, the results show that the images produced by the DHM system can be exploited to expand monitoring capabilities. Exploratory analyses illustrate the potential for anticipating viability decline and predicting IgG titer. These findings highlight the value of DHM for continuous culture monitoring in bioproduction environments. Future work will focus on integrating the setup into on-line and in-line configuration as well as validating our exploratory results.

## Materials and Methods

### A. Details on the optical set-up

DHM is implemented through a defocus microscope, using a custom-built inverted microscope^44^, equipped with a 0.25 NA air objective (Olympus PLN10X) matched with a 100 mm focal length infinity-corrected tube lens (Thorlabs TTL-100A), yielding an optical magnification of 5.56. Images were captured with a monochrome camera (IDS UI-3880-SE) equipped with a 3088 x 2076 pixel CMOS sensor of pixel pitch 2.4 *µ*m, yielding 0.432 *µ*m/pixel in the object plane after magnification, and a field of view of 1.17 mm^2^. Illumination was provided by a blue LED emitter, through a 400 *µ*m pinhole, and spectrally filtered at 450 nm with 10 nm FWHM. The optical setup includes a motorized stage for automated sample scanning (Thorlabs MLS203-1, controlled by a Thorlabs BBD302 controller) and a piezoelectric actuator (PiezoConcept FOC-300) to precisely position the objective at the desired distance from the sample.

To reproducibly achieve a fixed defocus, the sample plane is first located by acquiring a stack of 31 images spaced 10 *µ*m apart and computing the pixel-value standard deviation for each image. The focal plane is identified as the image with the lowest standard deviation, and the objective is translated by the specified offset to the desired defocus distance. For this type of images, minimal standard deviation of the image pixel values is a reliable indicator of the focal plane as long as distance to the focal plane is within reasonable limits (from tens to a few hundred *µ*m). Indeed, at focus, cells appear nearly transparent in transmission images, leading to minimal intensity variation and therefore a low standard deviation. This criterion was validated against manual focusing and proved robust. Particular attention was paid to the choice of defocus distance, as this parameter directly affects holographic reconstruction quality. Increasing defocus enhances the concentric interference fringes around each cell (observable in the Figure 1B), improving phase retrieval accuracy^45^. However, at high cell densities, excessive defocus causes signal overlap and complex interference patterns that degrade reconstruction. A relatively low defocus distance of 50 *µ*m was therefore chosen to maitain satisfactory reconstruction quality even at high cell densities.

The slide holder used in this study could accommodate four microscope slides, each containing four chambers, for a total of 16 independent counting chambers. Four images were taken per chamber, and each culture condition was represented in at least two chambers, yielding a minimum of eight images per condition. This number was found sufficient to capture enough cells for reliable viability analysis. The bench also enables bright-field imaging, which was used only during development for visual inspection of the cells and exploratory identification of relevant features.

### B. Details on the Image Reconstruction algorithm

Phase and absorption images were reconstructed using an updated version of the algorithm developed by Hervé et al^37^. In their work, the neural network was trained on simulated datasets for both phase and absorption reconstruction. In the present study, two key modifications were introduced. First, two separate models were trained, one dedicated to phase image prediction and another to absorption image prediction, as absorption reconstruction was unsatisfactory in the original approach. For phase reconstruction, the strategy of training on simulated data was retained, but substantially improved by incorporating greater variability in the morphology and spatial organization of the simulated cells. For absorption, the neural network was trained on an experimental dataset composed of 5,000 pairs of holograms and corresponding bright-field images, with the latter serving as the ground truth. The network takes as input not only a preliminary estimate of the sample but also the phase image obtained from the phase image prediction step.

### C. Details on the segmentation process

Cellpose^38^ is a deep learning-based, general-purpose cell segmentation algorithm. It employs a convolutional neural network trained on multiple biological image datasets. The network predicts spatial gradients (flow fields) that point from each pixel to the center of its corresponding cell, enabling accurate mask reconstruction even in crowded or irregularly shaped cells. Cellpose is widely used in the field of biological image analysis due to its strong performance and the ability to fine-tune the model on specific datasets. For this study, 20 images spanning a wide range of viability states and cell concentrations was semi-manually annotated and used to fine-tune the network. Performance was then evaluated on 16 independent images. Segmentation scores, assessed against manual annotations, consistently exceeded 98%, demonstrating the robustness of the algorithm for our application.

### D. Details on Data curation

Several filters were applied to remove incorrect cell detections from the analysis. Errors may arise either from the reconstruction algorithm, particularly hallucinations introduced by the neural network, or from the segmentation step. To limit these effects, detected objects exhibiting abnormally small diameters or implausible OPD or absorption values were excluded. Time points were also discarded from the study. A first reason was an insufficient number of detected cells, typically at the beginning of a culture or when an acquisition error occurred as A minimum number of cells, set to 1000 was required to obtain a representative histogram for the viability algorithm. Secondly, time points corresponding to clearly anomalous Vi-CELL viability measurements were removed after validation from biologists.

### E. Details on the viability measurement algorithm

At each time point, a 2D histogram was constructed using two features: mean OPD over the cell area and the mean absorption around the cell’s center of mass. Minimum and maximum histogram values were fixed across all time points and experiments. This histogram was treated as an image, and serves as the input for the viability analysis. A Gaussian filter (with a sigma of 3 pixels) was first applied to smooth small variations. A local maxima detection algorithm was then used to identify distinct cell populations within the culture. When multiple peaks are detected, only those exceeding a specified intensity threshold and separated by at least 8 pixels from neighboring peaks were retained. If a single peak was detected, the histogram image was passed through a neural network to determine whether the population was primarily composed of viable or non-viable cells. Depending on the population composition, the decision boundary was placed differently: if the cells was mainly alive, the line passes through a point below and to the left of the peak; if the population was a mixture of viable and non-viable cells, the line passes between the two peaks, closer to the right one; and if the population was mainly composed of non-viable cells, the line passes to the right of the peak. The line angle was constant in all cases. All parameters, including the angle, the distance from the peak for each population type, and the minimum distance between peaks, were determined through a combination of manual design and automatic optimization.

### F. Details on training and network implementation of the CNN used for the viability measurement algorithm

An EfficientNet V2-S^46^ model was employed to infer the population type from histogram images. A one-versus-all training and testing strategy ensured sufficient variability in the training set while enabling independent evaluation across multiple cultures. The network was trained using the ADAM optimizer with a binary cross-entropy (BCE) loss. A sampling strategy was applied to maintain equal proportions of each label in the training set. An early stopping procedure based on the training loss was used to prevent overfitting. The model takes as input the 2D OPD/absorption histogram, along with nine additional 2D histograms generated from combinations of the following eight features: cell diameter, eccentricity, area, granularity at two levels corresponding to very fine structures (approximately the size of a pixel, i.e., 0.432 *µ*m), optical volume difference (OVD, defined as the integrated optical path difference across the cell, which is directly related to cell dry mass), and both the standard deviation and median absolute deviation of the OPD across the cell surface. Granularity was computed following the method proposed in^47^

### G. Details on the UMAP procedure

UMAP (Uniform Manifold Approximation and Projection)^40^ is a dimensionality reduction technique that projects high-dimensional data into lower dimensions, typically 2D, while preserving both local and global structure. This makes patterns, clusters, and relationships in complex datasets easier to interpret. In this study, UMAP was employed to integrate multiple cellular features into a 2D visualization, facilitating intuitive exploration and identification of population dynamics. The ten features used in the viability measurement algorithm were analyzed with the following UMAP parameters: euclidean distance as the metric, 100 neighbors, and a minimum distance of 0.8. These parameter values were chosen empirically. Several combinations were tested, and the corresponding projection that appeared most informative was selected.

### H. Details on the IgG titer prediction

The same features used for the UMAP projection were employed to predict the IgG titer. Time points with viability above 85% were selected, as later points are strongly affected by the presence of non-viable cells, and harvest typically occurs before viability drops too low. Prediction was performed at the time-point level: all cell features were aggregated into 10-bin histograms, yielding 10 values per feature and 120 input features in total. This approach was chosen over simpler statistics, such as the first moment, because histograms better capture the intrinsic shape of the distribution. Several regression algorithms were evaluated, including linear regression, random forest, Lasso, Ridge, and SVR. Gradient boosting regression was selected for its slightly superior performance. As this analysis was intended for demonstration purposes, parameters were not optimized and default settings were used.

### i. Cell line description

Cell line C-A.1 corresponds to a CHO-DG44 cell line. Cell line C-A.2 corresponds to a CHO-K1 cell line. Cell line C-B corresponds to a CHO-K1–derived cell line. Cell line C-C1 corresponds to a CHO-DXB11–derived cell line. Cell line C-C2 corresponds to a CHO-K1–derived cell line. Cell line C-D corresponds to a proprietary GS-CHO cell line.

## Acknowledgements

The authors warmly thank Geoffrey Esteban, Iprasense France for his coordination of the SELPHi project, which led to this publication. The authors would also like to thank Cedric Allier for initiating the study, and Caroline Paulus, Ondrej Mandula, Olivier Cioni, Vincent Remondière and Xavier Mermet for for their valuable discussions, technical assistance, and constructive feedback throughout this study.

## Author contributions statement

G.Godefroy, S.Lhomme, E.Guedon and E.Calvosa conceived the study.

T.Cantat-Moltrech developed the DHM acquisition setup.

G.Godefroy, L.Hervé and G.Girard developed the image reconstruction.

G.Godefroy and G.Girard developed the image analysis and viability estimation algorithms.

A.Berger, T.Saillard and A.La and performed the experiments.

G.Godefroy drafted the manuscript with input from all authors.

All authors approved the final version of the manuscript.

## Data availability

The datasets generated during the current study are not publicly available due to industrial confidentiality constraints but are available from the corresponding author on reasonable request and subject to approval by the data owners.

## Ethics Declarations

The experiments were conducted using established Chinese hamster ovary (CHO) cell lines and derivatives, including CHO-K1, CHO-DG44, CHO-DXB11, proprietary GS-CHO, and CHO-K1–derived cell lines. As these are immortalized cell lines and the study did not involve human participants or live animals, ethical approval was not required.

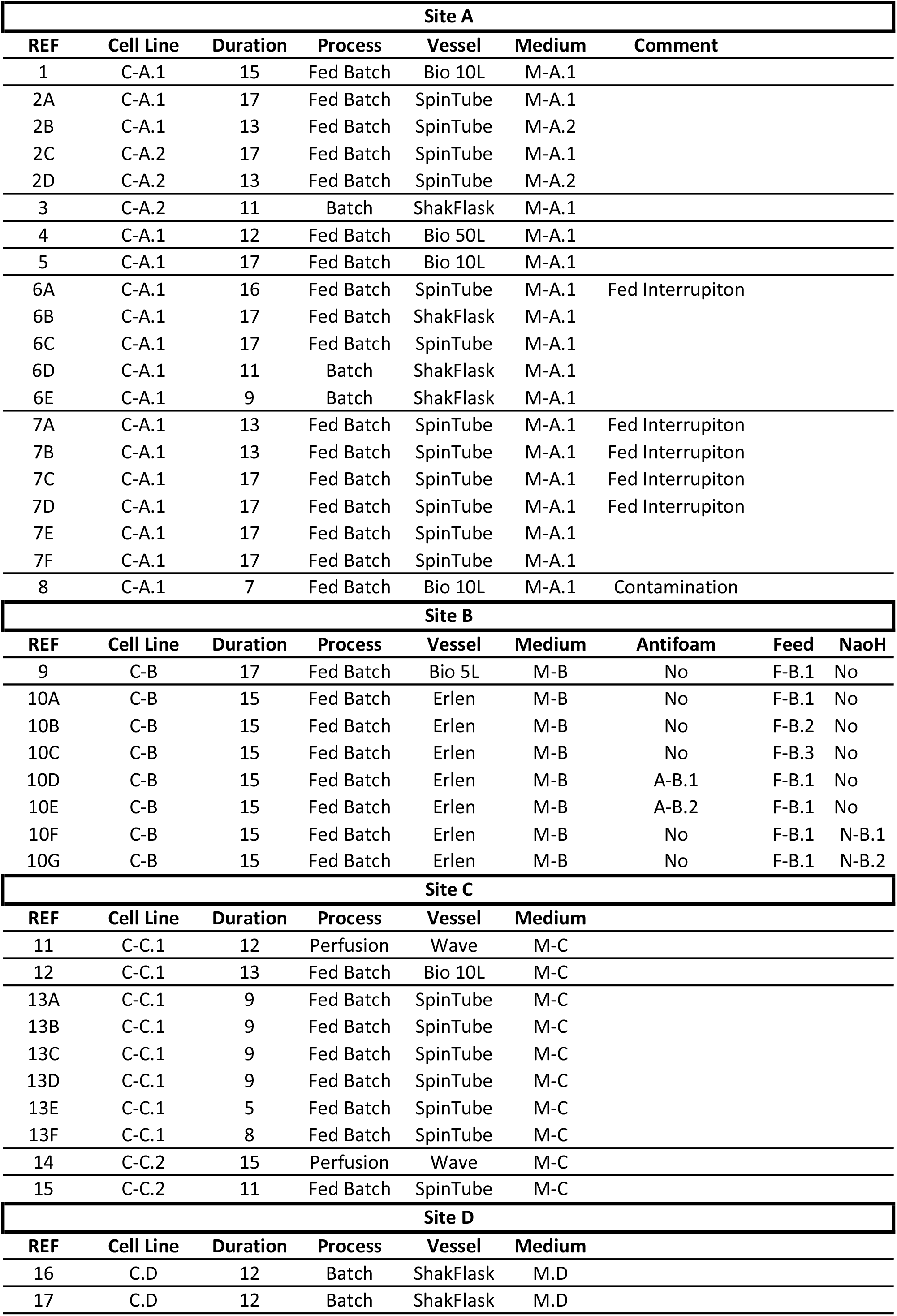

